# Genome-Wide Mapping In A House Mouse Hybrid Zone Reveals Hybrid Sterility Loci And Dobzhansky-Muller Interactions

**DOI:** 10.1101/009373

**Authors:** Leslie M. Turner, Bettina Harr

## Abstract

Mapping hybrid defects in contact zones between incipient species can identify genomic regions contributing to reproductive isolation and reveal genetic mechanisms of speciation. The house mouse features a rare combination of sophisticated genetic tools and natural hybrid zones between subspecies. Male hybrids often show reduced fertility, a common reproductive barrier between incipient species. Laboratory crosses have identified sterility loci, but each encompasses hundreds of genes. We map genetic determinants of testis weight and testis gene expression using offspring of mice captured in a hybrid zone between *M. musculus musculus* and *M. m. domesticus*. Many generations of admixture enables high-resolution mapping of loci contributing to these sterility-related phenotypes. We identify complex interactions among sterility loci, suggesting multiple, non-independent genetic incompatibilities contribute to barriers to gene flow in the hybrid zone.

## INTRODUCTION

New species arise when reproductive barriers form, preventing gene flow between populations (Coyne and Orr 2004). Recently, two approaches have substantially advanced understanding of the genetic mechanisms underlying reproductive isolation (reviewed in Noor and Feder 2006; reviewed in Wolf et al. 2010). Genetic crosses in the laboratory involving model organisms have identified loci and genes causing hybrid defects, a common type of reproductive barrier caused by genetic interactions between divergent alleles (Bateson 1909; Dobzhansky 1937; Muller 1942). In nature, recent technological advances enable fine-scale characterization of genome-wide patterns of divergence between incipient species and variation in hybrid zones.

For example, ‘islands of divergence’ have been reported in species pairs from taxonomically diverse groups (Turner et al. 2005; Nadeau et al. 2011; Nosil et al. 2012; Ellegren et al. 2013; Hemmer-Hansen et al. 2013; Renaut et al. 2013; Carneiro et al. 2014; Poelstra et al. 2014; Schumer et al. 2014). These high-divergence genomic outlier regions are sometimes referred to as ‘islands of speciation,’ resistant to introgression because they harbor genes causing reproductive isolation. However, other forces can create similar genomic patterns, thus islands may not always represent targets of selection that contributed to speciation (Noor and Bennett 2009; Turner and Hahn 2010; Renaut et al. 2013; Cruickshank and Hahn 2014).

An alternative approach to identify genomic regions contributing to reproductive isolation is to map known reproductive barrier traits in naturally hybridizing populations. The potential for mapping in hybrid zones is long-recognized (Kocher and Sage 1986; Harrison 1990; Szymura and Barton 1991; Briscoe et al. 1994; reviewed in Rieseberg and Buerkle 2002). Hybrid zones are “natural laboratories for evolutionary studies” (Hewitt 1988) enabling investigation of speciation in progress. The Dobzhansky-Muller model predicts that hybrid incompatibilities between incipient species accumulate faster than linearly with time (Orr 1995), thus investigating taxa in the early stages of speciation facilitates identification of incompatibilities that initially caused reproductive isolation vs. incompatibilities that arose after isolation was complete.

Despite these advantages, few studies have mapped barrier traits or other fitness-related traits in nature, due to the logistical challenges of collecting dense genome-wide genetic markers in species with admixed populations and well-characterized phenotypes. Examples include associations between pollen sterility and genomic regions showing reduced introgression in a sunflower hybrid zone (Rieseberg et al. 1999) and loci contributing to variation in male nuptial color and body shape mapped in a recently admixed stickleback population (Malek et al. 2012).

House mice (*Mus musculus*) are a promising model system for genetic mapping in natural populations (Laurie et al. 2007), and have an abundance of genetic tools available to ultimately isolate and characterize the causative genes underlying candidate loci. Three house mouse subspecies - *M. m. musculus, M. m. domesticus* and *M. m. castaneus* - diverged ~500,000 years ago from a common ancestor (reviewed in Boursot et al. 1993; Salcedo et al. 2007; Geraldes et al. 2008). *M. m. musculus* and *M. m. domesticus* (hereafter *musculus* and *domesticus*) colonized Europe through different geographic routes and meet in a narrow secondary contact zone running through central Europe from Bulgaria to Denmark (Sage et al. 1986; Boursot et al. 1993). Genome-wide analyses of patterns of gene-flow in several geographically distinct transects across the hybrid zone have identified genomic regions showing reduced introgression, which may contribute to reproductive isolation (Tucker et al. 1992; Macholan et al. 2007; Teeter et al. 2008; 2010; Janousek et al. 2012).

Reduced male fertility is common in wild-caught hybrids (Turner et al. 2012; Albrechtová et al. 2012) and in *musculus* - *domesticus* hybrids generated in the laboratory (Britton-Davidian et al. 2005; reviewed in Good et al. 2008a), implying hybrid sterility is an important barrier to gene flow in house mice. Mapping studies using F_1_, F_2_ and backcross hybrids generated from laboratory crosses between house mouse subspecies have identified many loci and genetic interactions contributing to sterility phenotypes (Storchova et al. 2004; Good et al. 2008b; White et al. 2011; Dzur-Gejdosova et al. 2012; Turner et al. 2014). *Prdm9*, a histone methyltransferase, was recently identified as a gene causing F_1_ hybrid sterility, and is the first mammalian hybrid incompatibility gene identified (Mihola et al. 2009). Comparisons between different F_1_ crosses show that hybrid sterility alleles are polymorphic within subspecies (Britton-Davidian et al. 2005; Good et al. 2008a). Furthermore, reduced fertility phenotypes observed in nature vary in severity; complete sterility, as documented in some F_1_ crosses, appears to be rare or absent in the hybrid zone (Turner et al. 2012; Albrechtová et al. 2012). Taken together, studies of hybrid sterility in house mice indicate that, even in the early stages of speciation, the genetic basis of hybrid defects can be complex. Studies of gene flow in the hybrid zone and of hybrid sterility in the laboratory both have advantages and have shed light on the speciation process. Mapping sterility phenotypes in natural hybrids can potentially integrate insights from the two approaches by identifying associations between hybrid incompatibility loci and reduced gene flow across the hybrid zone.

Here, we map sterility-related phenotypes in hybrid zone mice to investigate the genetic architecture of reproductive isolation between incipient species. We performed a genome-wide association study (GWAS) to map testis weight and testis gene expression in 185 first generation lab-bred offspring of wild-caught hybrid mice (Figure 1-figure supplement 1). GWAS have been powerful in humans, loci contributing to hundreds of quantitative traits associated with disease and other phenotypic variation have been identified (reviewed in Stranger et al. 2011). Examples of GWAS for fitness-related traits in non-humans are only beginning to emerge (Johnston et al. 2011; Filiault and Maloof 2012; Magwire et al. 2012).

Our hybrid zone GWAS identified genomic regions associated with variation in relative testis weight (testis weight/body weight) and genome-wide testis expression pattern, including regions previously implicated in hybrid sterility as well as novel loci. Motivated by the Dobzhansky-Muller genetic model of hybrid defects, we tested for genetic interactions (Dobzhansky-Muller interactions – “DMIs”) between loci affecting testis weight or expression pattern. All loci except one showed evidence for interaction with at least one partner locus, and most interact with more than one partner. The deviations in phenotype associated with most interactions were large - affected individuals have phenotypes below the range observed in pure subspecies – suggesting these interactions indeed are hybrid incompatibilities. To our knowledge, this is the first GWAS for a reproductive barrier trait. Using natural hybrids provided high mapping resolution that will facilitate future studies to identify causative genes; for example, a majority of GWAS regions contain 10 or fewer genes. Moreover, this study provides the first genome-scale description of a hybrid incompatibility network in nature.

## RESULTS

### Sterility-associated phenotypes

We investigated two phenotypes in males from the house mouse hybrid zone: relative testis weight (testis weight/ body weight) and genome-wide testis gene expression pattern. Both of these phenotypes have previously been linked to hybrid male sterility in studies of mice from crosses between *musculus* and *domesticus* and mice from the hybrid zone (Britton-Davidian et al. 2005; Rottscheidt and Harr 2007; reviewed in Good et al. 2008a; 2010; White et al. 2011; Turner et al. 2012; 2014). We refer to these as “sterility phenotypes,” following conventional terminology in the field, however, it is important to note that the severity of defects observed in most hybrid zone mice are consistent with reduced fertility/partial sterility (Turner et al. 2012).

Testis expression PC1 (explaining 14.6% of the variance) is significantly correlated with relative testis weight (cor = 0.67, *P* = 2x10^-16^) indicating there is a strong association between those two sterility phenotypes (Figure 1 - figure supplement 2). Principal component 2 (PC2, 8.1% variance) is strongly correlated with hybrid index (% *musculus* autosomal SNPs: cor = 0.75, *P* = 2x10^-16^), thus the effect of hybrid defects do not obscure subspecies differences in expression.

In the mapping population, 19/185 (10.2%) individuals had relative testis weight below the minimum observed in pure subspecies males and 21/179 (11.7%) individuals had expression PC1 scores below (PC1 = −46.97) the pure subspecies range.

### Association mapping

We identified four SNPs significantly associated with relative testis weight in three regions on the X chromosome using stringent threshold determined by permutation (Table 1; Figure 1A). An additional 51 SNPs were significant using a more permissive significance threshold (false discovery rate (FDR)<0.1). Significant SNPs were clustered in 12 genomic regions (of size 1 bp – 13.3 Mb; Table 1). We report GWAS regions defined using the permissive FDR threshold because we plan to combine mapping results from multiple phenotypes to identify candidate sterility loci, based on the idea that spurious associations are unlikely to be shared among phenotypes. Significant regions were located on the X chromosome and 9 autosomes, suggesting a minimum of 10 loci contribute to variation in testis weight. It is difficult to estimate the precise number of genes involved, because the extent of linkage disequilibrium (LD) around a causative mutation depends on the phenotypic effect size, recombination rate, allele frequency, and local population structure. Multiple significant regions might be linked to a single causative mutation, or conversely, a significant region might be linked to multiple causative mutations in the same gene or in multiple genes.

**Figure 1.**
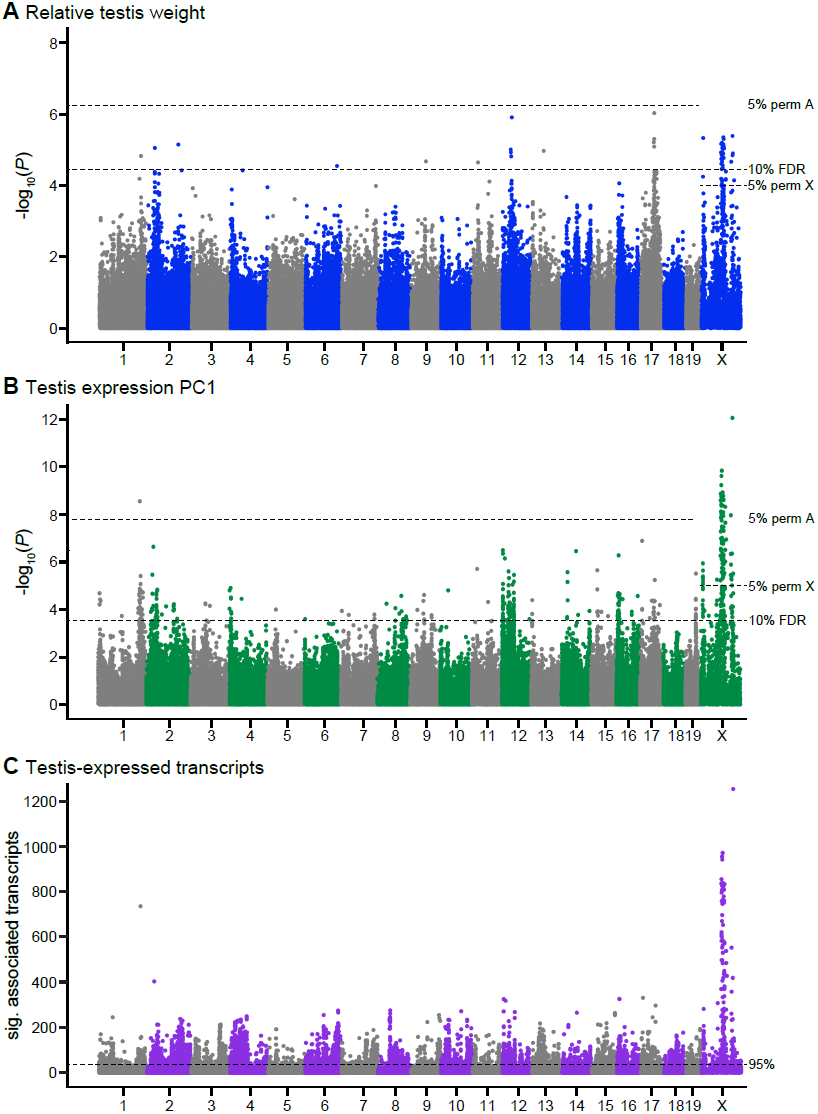
Single SNPs associated with (**A**) relative testis weight, (**B**) testis expression principal component 1, and (**C**) expression of transcripts located on other chromosomes (*trans*). Dashed lines indicate significance thresholds based on: permutations for autosomes (labeled 5% perm A), permutations for X chromosome (labeled 5% perm X), false discovery rate < 0.1 (labeled 10% FDR), and 95^th^ percentile of significant transcript association counts across SNPs (labeled 95%).

**Table 1.**
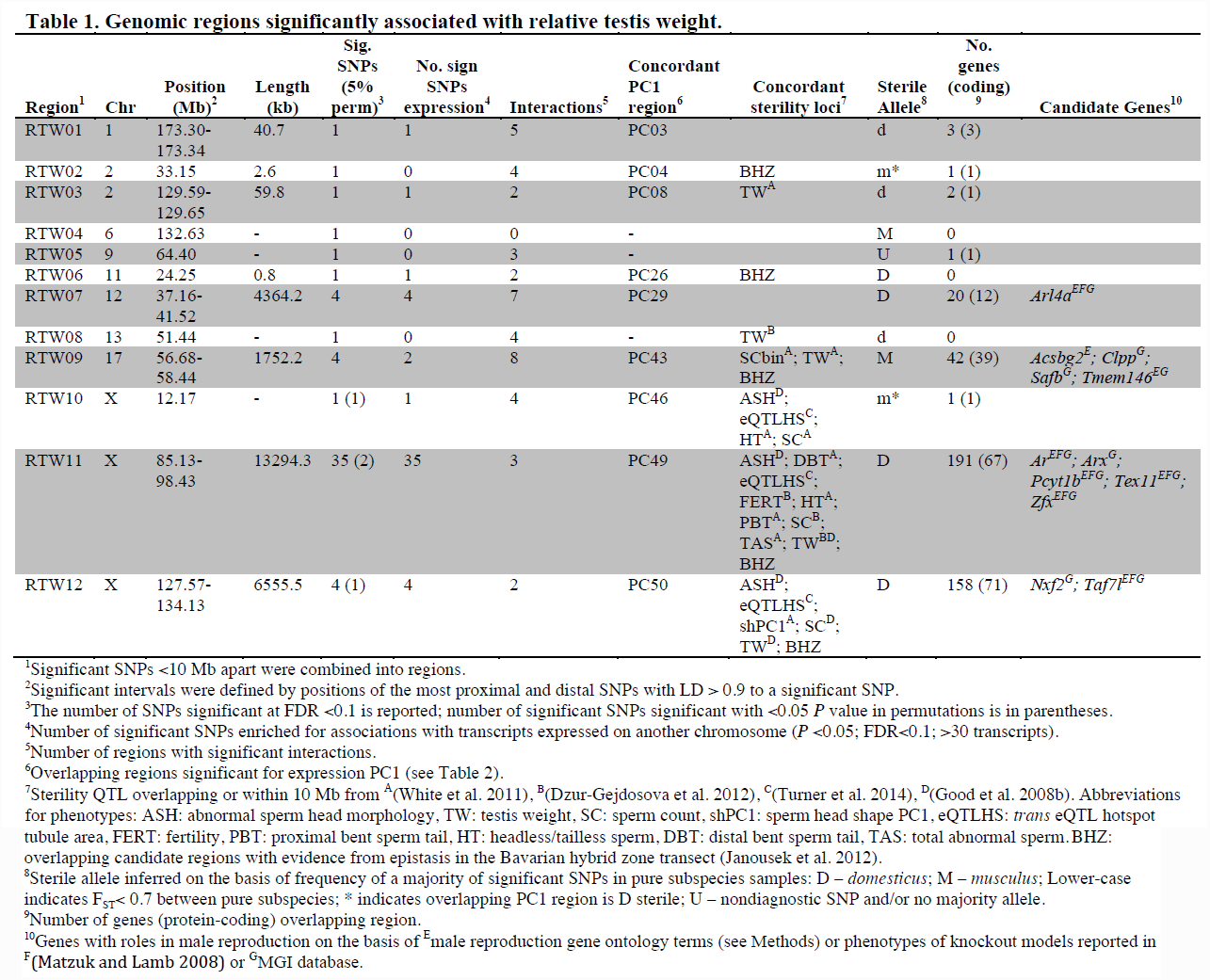
Genomic regions significantly associated with relative testis weight.

We identified 104 SNPs on the X and chromosome 1 significantly associated with expression PC1 using stringent permutation-based thresholds (Table 2; Figure 1B). An additional 349 SNPs were significant with the more permissive threshold of FDR <0.1. Significant SNPs clustered in 50 genomic regions located on 18 autosomes and the X.

**Table 2.**
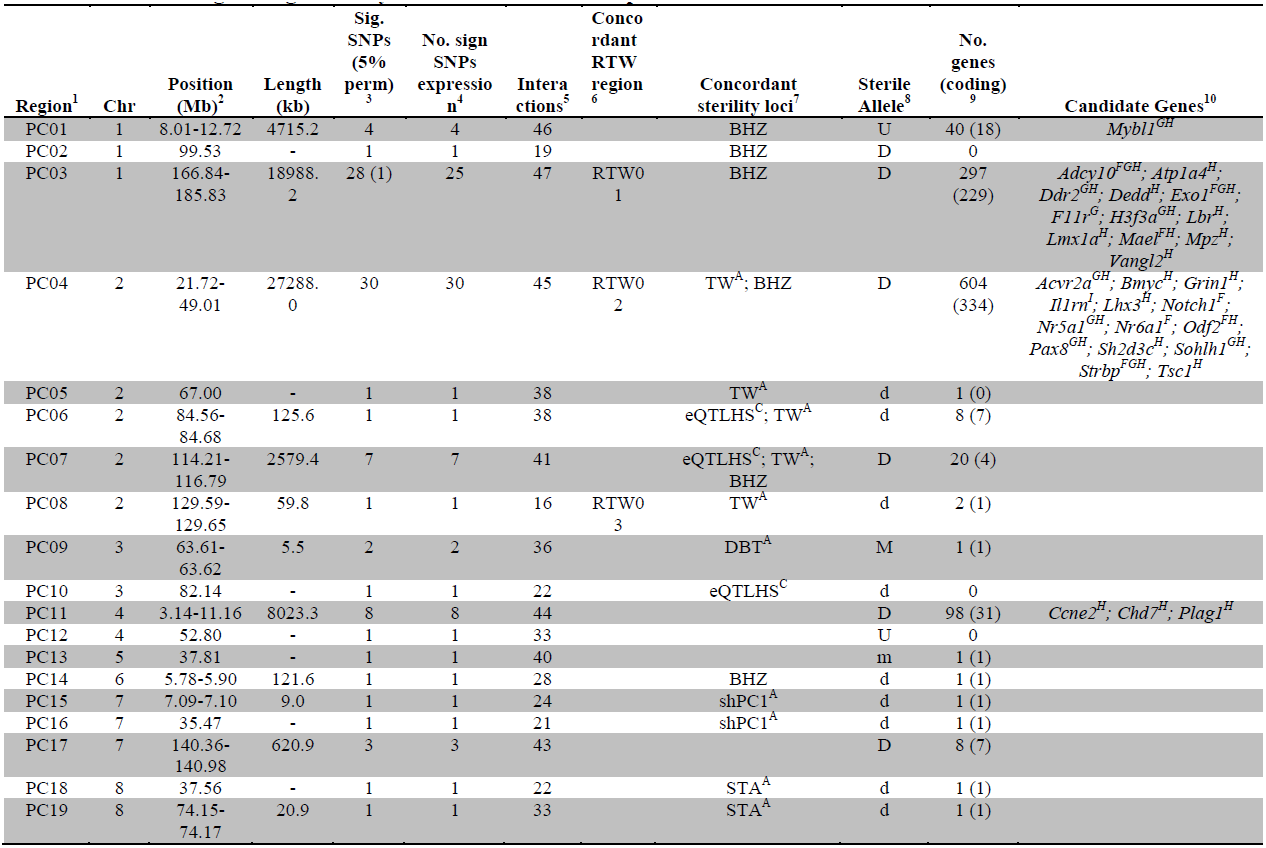
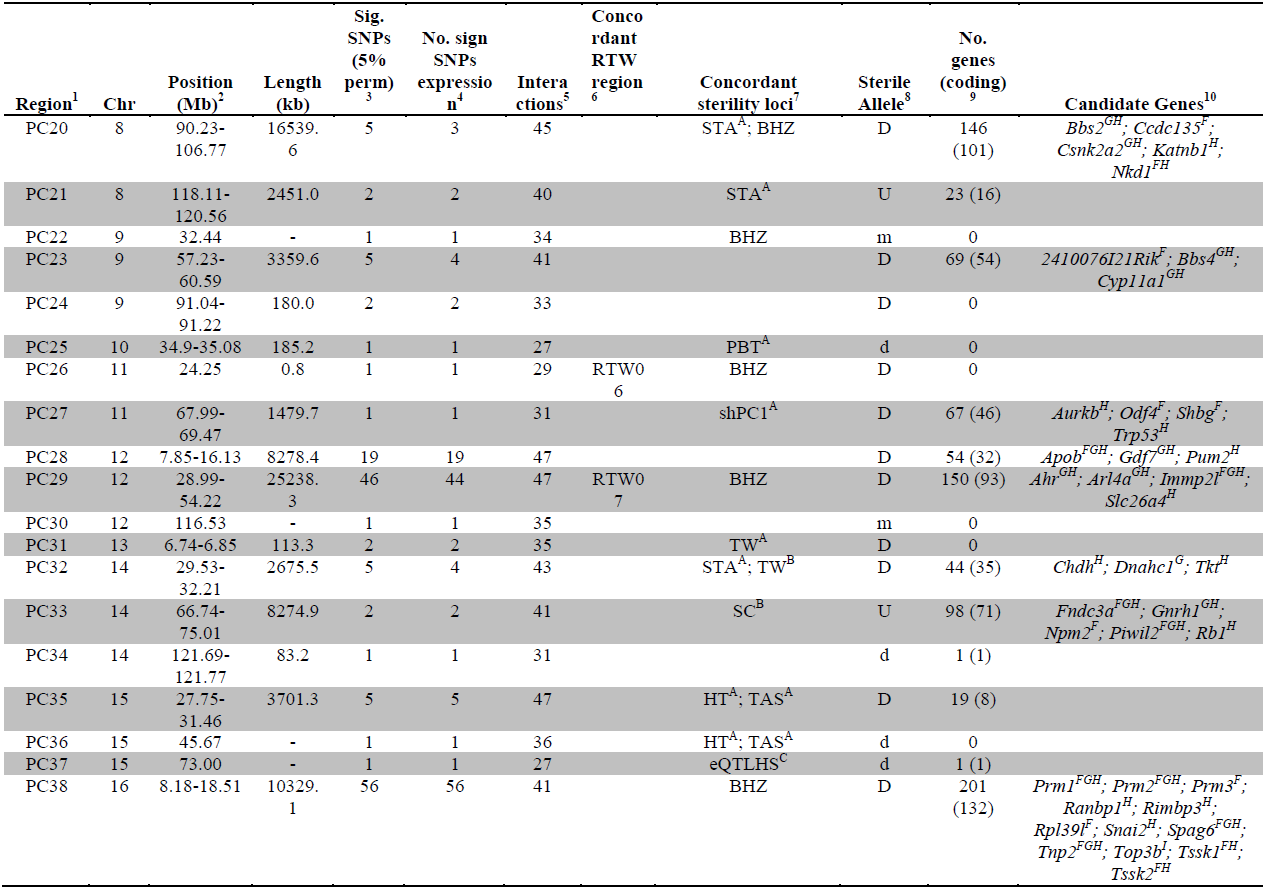
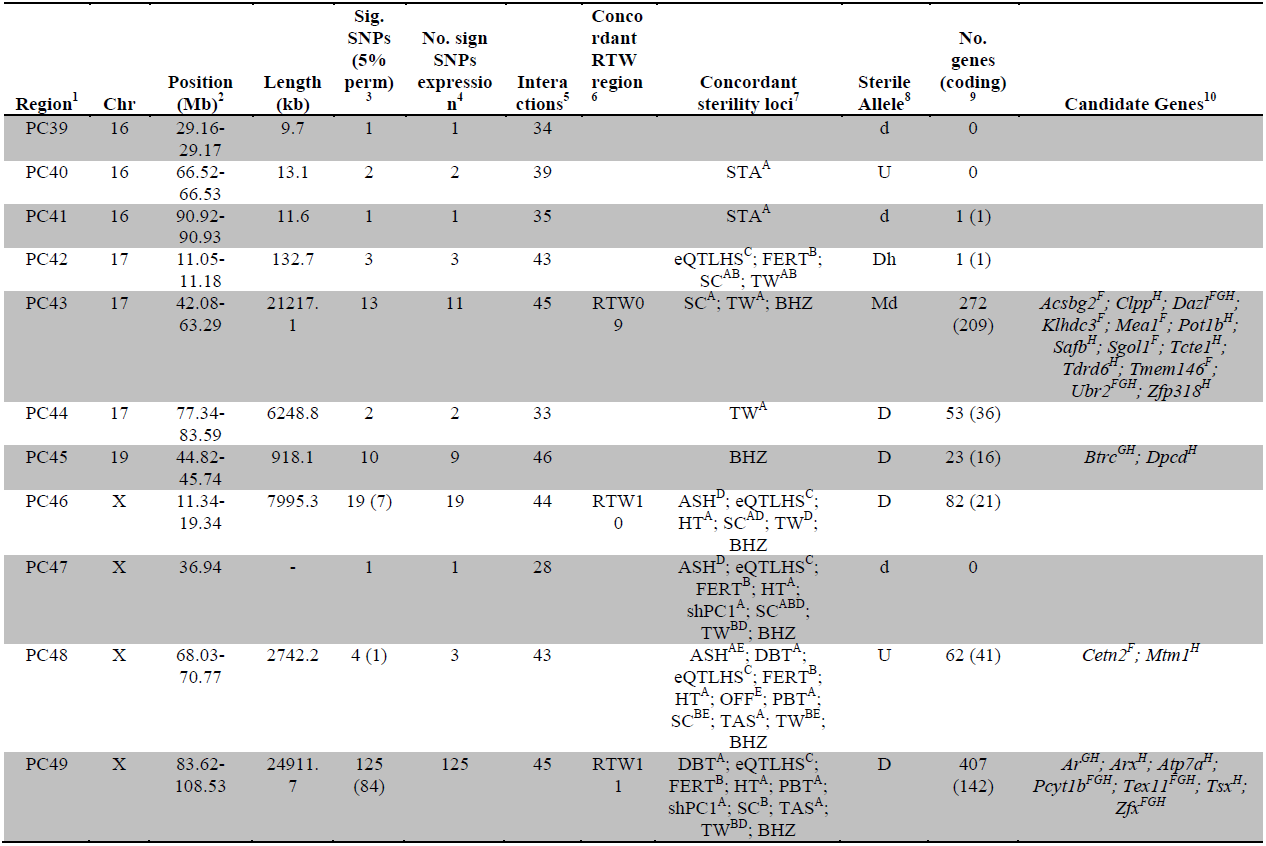
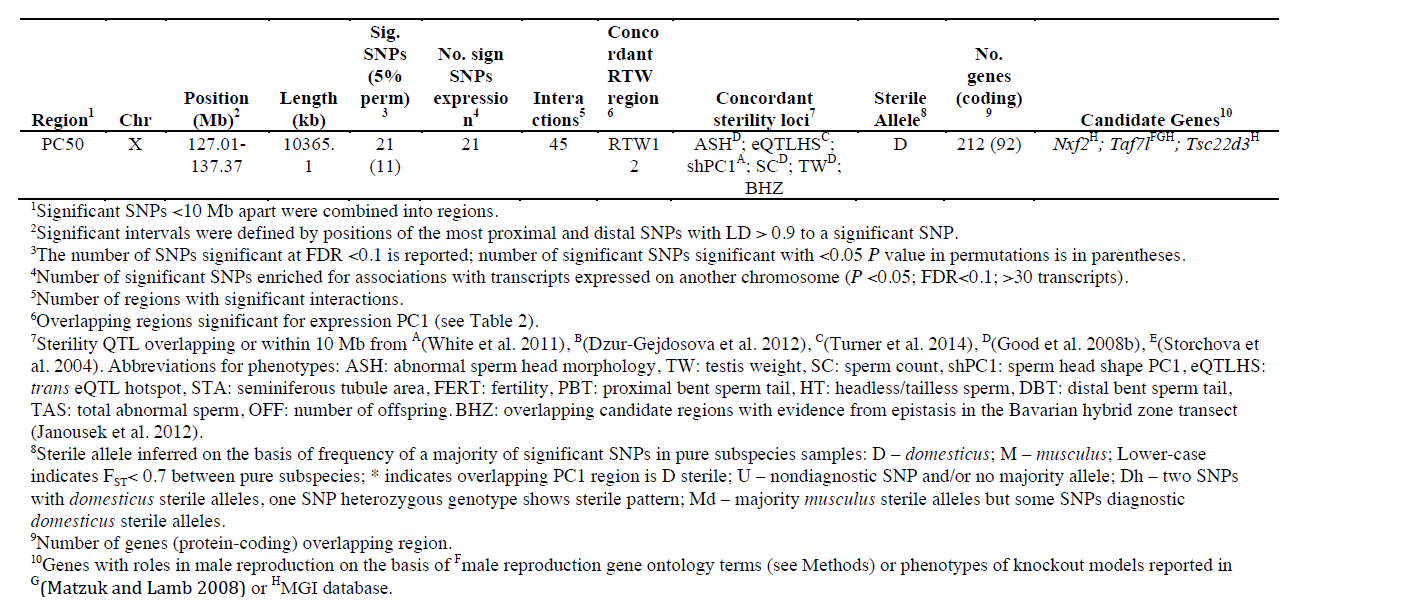
Genomic regions significantly associated with testis expression PC1.

To gain further insight into associations between sterility and gene expression, we mapped expression levels of individual transcripts. A total of 18,992/36,323 probes showed significant associations with at least on SNP. We focused on *trans* associations (SNP is located on different chromosome from transcript), based on evidence from a study in F_2_ hybrids that *trans* expression QTL (eQTL) are associated with sterility while *cis* eQTL are predominantly associated with subspecies differences (Turner et al. 2014). To identify SNPs significantly enriched for *trans* associations with expression, we used a threshold set to the 95% percentile counts of significantly associated probes across all SNPs (30 probes, Figure 1C).

There was substantial overlap between mapping results for testis weight and expression PC1; 48/55 SNPs significant for relative testis weight (9 regions) were also significant for expression PC1. A permutation test, performed by randomly shuffling the positions of GWAS regions in the genome, provides strong evidence that this overlap is non-random (*P*<0.0001, 10,000 permutations). Most SNPs significant for testis weight and/or expression PC1 were significantly enriched for *trans* associations with individual transcripts (relative testis weight: 49/55 SNPs, 8/12 regions; PC1: 440/453 SNPs, 50/50 regions). The combined mapping results provide multiple lines of evidence for contributions of all 50 PC1 regions and 9/12 testis weight regions. The three testis-weight regions (RTW04, RTW05, RTW08) not significantly associated with testis expression phenotypes are more likely to be spurious and are weaker candidates for future study.

### Genetic interactions

Power to identify pairwise epistasis in GWAS for quantitative traits is limited even with very large sample sizes, due to multiple testing issues (e.g. Marchini et al. 2005). The Dobzhansky-Muller predicts that the effect of each hybrid defect gene depends on interaction with at least one partner locus. Hence, for hybrid sterility traits, there is a hypothesis-driven framework in which to limit tests for epistasis to a small subset of possible interactions.

We tested for genetic interactions between all pairs of significant SNPs (FDR <0.1) located on different chromosomes for testis weight and for expression PC1. We identified 142 significant pairwise interactions for relative testis weight, representing 22 pairs of GWAS regions (Figure 2A). These results provide evidence for a minimum of 13 autosomal-autosomal and five X – autosomal interactions affecting testis weight.

**Figure 2.**
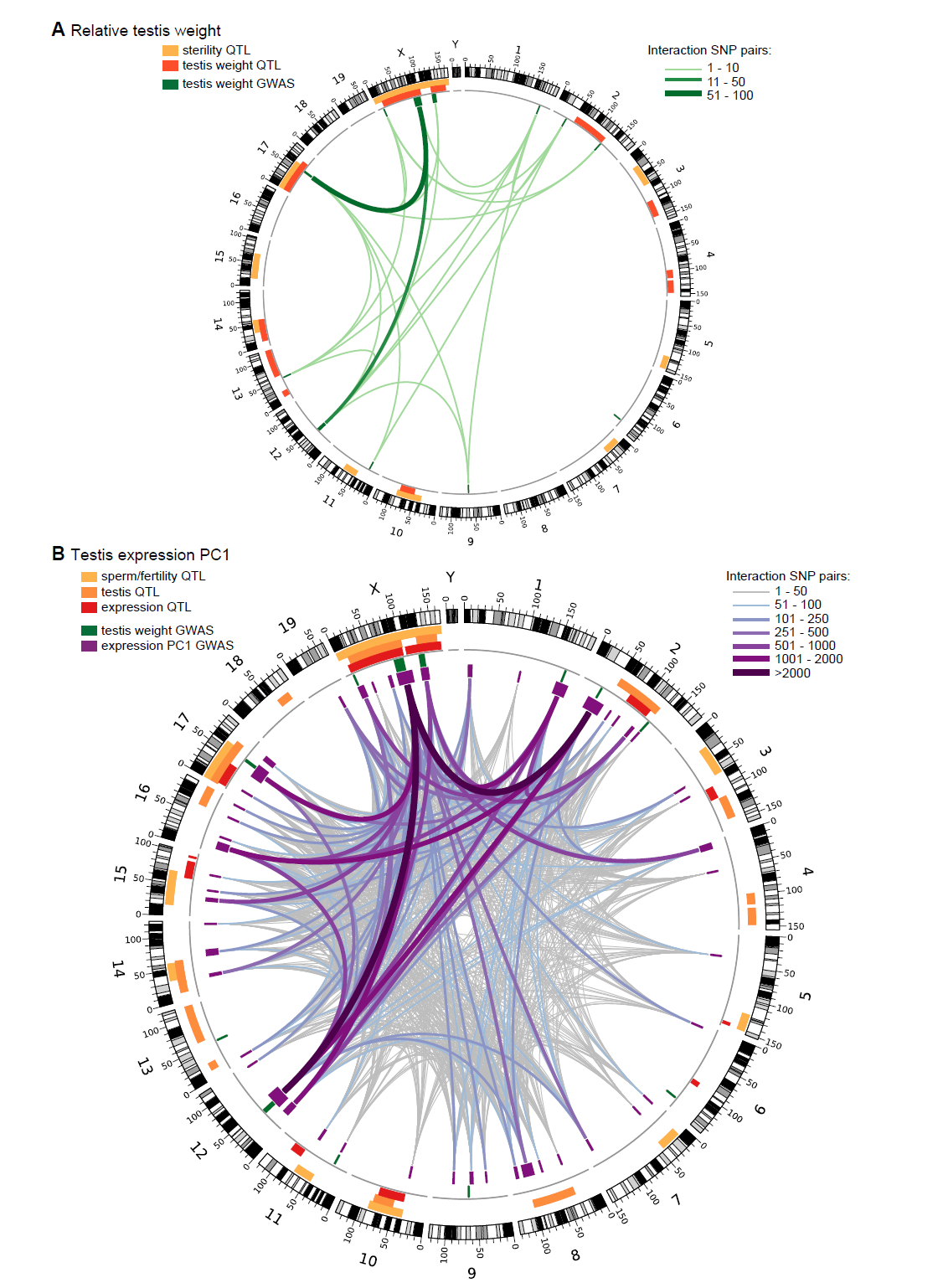
Significant GWAS regions and interactions associated with (**A**) relative testis weight and (**B**) testis expression principal component 1 in hybrid zone mice. In (**A**), orange and yellow boxes in outer rings (outside grey line) indicate quantitative trait loci (QTL) identified for testis weight and other sterility phenotypes in previous studies (see Table 1 for details). Green boxes indicate significant GWAS regions for relative testis weight. Green lines represent significant genetic interactions between regions; shade and line weight indicate the number of significant pairwise interactions between SNPs for each region pair. In (**B**), orange boxes in outer rings indicate QTL for testis-related phenotypes (testis weight and seminiferous tubule area) identified in previous studies, yellow boxes indicate QTL for other sterility phenotypes and red boxes indicate *trans* eQTL hotspots (see Table 2 for details). Green boxes indicate significant GWAS regions for relative testis weight. Purple boxes indicate significant GWAS regions for testis expression PC1. Lines represent significant genetic interactions between regions; color and line weight – as specified in legend - indicate the number of significant pairwise interactions between SNPs for each region pair. Plot generated using circos (Krzywinski et al. 2009).

We identified 44,145 significant interactions between SNPs for expression PC1. The 913 GWAS region pairs provide evidence that at least 144 autosomal-autosomal interactions and 18 X-autosomal interactions contribute to expression PC1 (Figure 2B).

### Effect size

We used deviations from population means for single SNPs and two-locus genotypes to estimate the phenotypic effects of GWAS regions and interactions (Figure 3A, B). As expected, interactions had greater effects, on average, than single loci for both phenotypes (relative testis weight: single locus mean = −1.81 mg/g, interaction mean = −4.07 mg/g; expression PC1: single locus mean = −81.51, interaction mean = −130.77). We provide examples of autosomal-autosomal and X-autosomal SNP pairs with significant interactions for each phenotype in Figure 3C. It is important to note that mean deviations are rough estimates of effect sizes, which don’t account for family structure.

**Figure 3.**
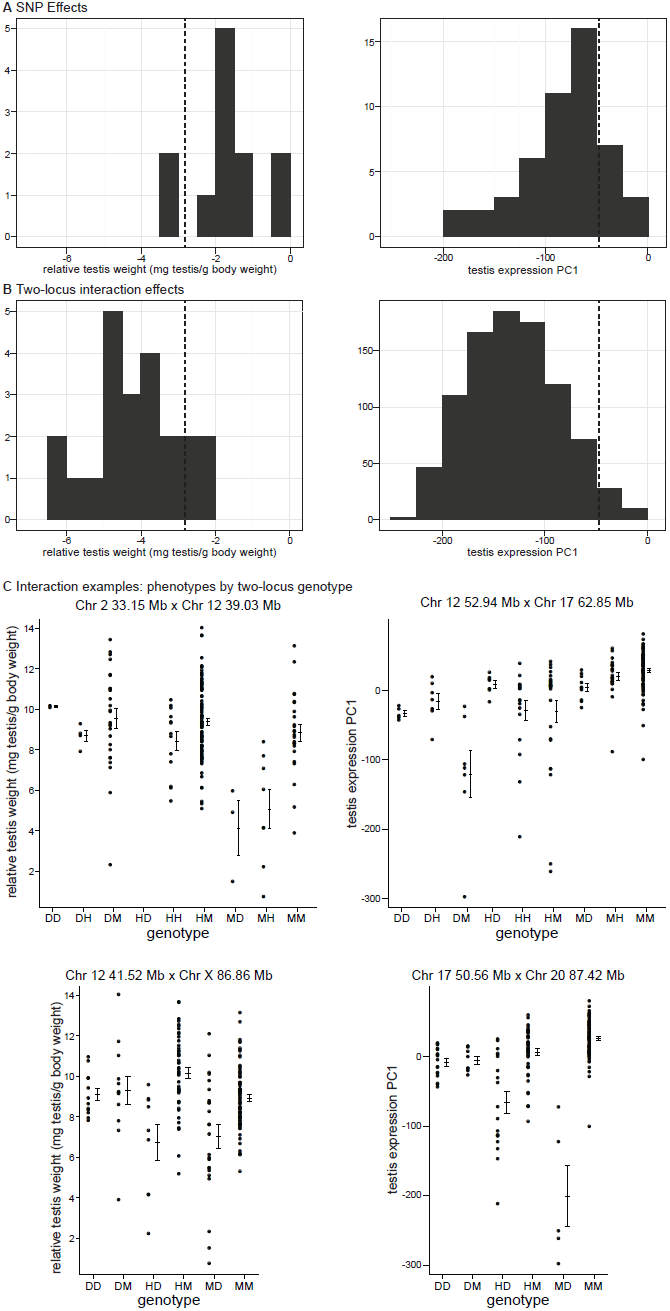
Phenotypic effects of testis-weight loci and interactions. Histograms showing maximum deviations from the population mean for (**A**) single SNPs and (**B**) two-locus interactions. Dashed vertical lines indicate minimum values observed in pure subspecies males. (**C**) Examples of phenotypic means by two-locus genotype for autosomal-autosomal and X-autosomal interactions. Genotypes are indicated by one letter for each locus: D – homozygous for the *domesticus* allele, H – heterozygous, M – homozygous *musculus*.

It is possible that some of the GWAS regions we mapped contribute to quantitative variation within/between subspecies, rather than hybrid defects. The lowest genotypic means for most interactions fell below the range observed in pure subspecies (relative testis weight: 19/22 (86.3%) region pairs; expression PC1: 877/913 (96%) region pairs; Figure 3A,B), consistent with the hypothesis that interactions represent Dobzhansky-Muller incompatibilities.

### Mapping simulations

We performed simulations to assess the performance of the mapping procedure for different genetic architectures by estimating the power to detect causative loci and the false positive rate. We simulated phenotypes based on two-locus genotypes from the SNP dataset using genetic models for nine genetic architecture classes (*i.e.* autosomal vs. X linked, varied dominance) with parameters based on the observed distribution of relative testis weight (Figure 4-figure supplements 1, Figure 4-Source data 1).

**Figure 4.**
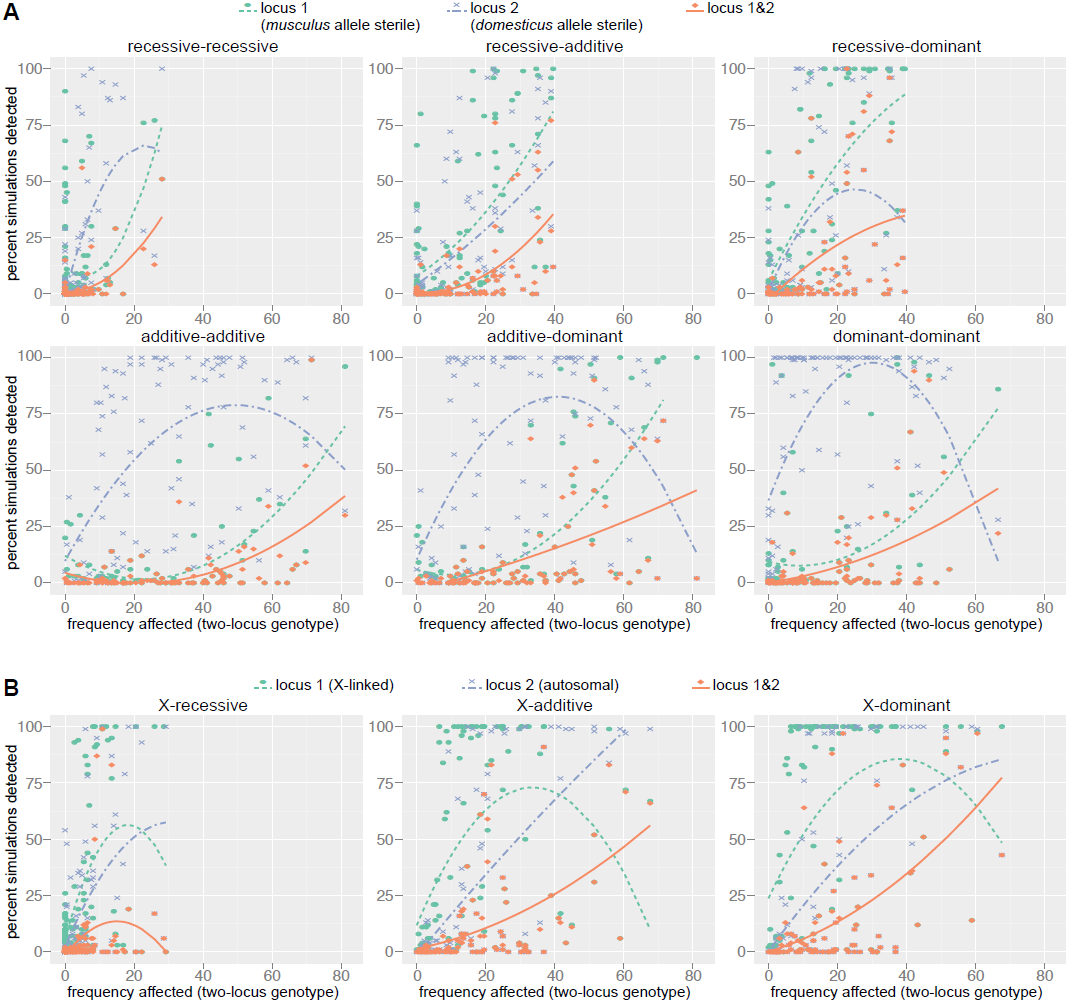
Mapping power in simulations. Each panel results from a single genetic architecture model for (**A**) 100 autosomal-autosomal SNP pairs and (**B**) 100 X-autosomal SNP pairs. Each point represents the percentage of data sets generated from a single SNP pair in which locus 1 (*domesticus* sterile allele; green), locus 2 (*musculus* sterile allele; purple), or both loci (orange) were identified by association mapping (≥1 SNP significant by permutation based threshold within 10 Mb of ‘causal’ SNP). The x axis indicates the percentage of individuals with partial or full sterility phenotypes. Curves were fit using 2^nd^ order polynomials. In (**A**), locus 1 indicates the SNPs with *musculus* alleles sterile and locus 2 indicates the SNPs with *domesticus* alleles sterile. In (**B**), locus 1 is the X-linked SNP and locus 2 is the autosomal SNP.

The distribution of distances to the causal SNP for all significant SNPs located on the same chromosome (Figure 4-figure supplement 2) shows that the majority of significant SNPs (62.7%), are within 10 Mb of the causal SNP, however a small proportion of significant SNPs are >50 Mb from the causal SNP. In most cases, causal SNPs detected at long distances also had significant SNPs nearby, for example 83.4% of loci with significant SNPs 1 – 10 Mb distant also has significant SNPs within 1 Mb. These results provide support for our choice to define significant GWAS regions by combining significant SNPs within 10 Mb, and suggest these regions are likely to encompass the causative gene.

As expected, the power to detect one or both causative loci depended on the location (autosomal vs. X-linked), dominance, and frequency of both ‘causative’ alleles (Table 3). For example, the mean percentage of simulations for which both loci were detected (SNP <10 Mb significant by permutation-based threshold) was six times higher (14.4%) for the X chromosome x autosomal dominant architecture compared to the autosomal-recessive x autosomal-recessive architecture (2.6%). The relationship between power and the proportion of affected individuals in the mapping population was complex. Interestingly, power was high for some simulations with very few affected individuals. In these cases, the few individuals carrying the lower frequency sterility allele by chance also carried the sterility allele from the second locus, thus the average effect was not diminished by individuals carrying one but not both interacting sterility alleles.

**Table 3.** Results of mapping simulations.

| Architecture<sup>1</sup> | Med. Distance to Causal SNP (Mb) X chromosome/Autosome | Locus 1 detected<sup>2,3</sup> |  |  | Locus 2 detected<sup>3,4</sup> |  |  | Both loci detected<sup>3</sup> |  |  | Mean No. Sig. SNPs |  |  |
| --- | --- | --- | --- | --- | --- | --- | --- | --- | --- | --- | --- | --- | --- |
|  |  | 0.2 Mb | 1 Mb | 10 Mb | 0.2 Mb | 1 Mb | 10 Mb | 0.2 Mb | 1 Mb | 10 Mb | 10 Mb<sup>5</sup> | 50 Mb<sup>5</sup> | Diff. Chr.<sup>6</sup> |
| Permutation $P<0.05$ | | | | | | | | | | | | | |
| rec-rec | 5.9 | 7.2 | 8.4 | 12.3 | 9.0 | 11.7 | 15.8 | 0.3 | 0.7 | 2.6 | 1.1 | 1.5 | 5.5 |
| rec-add | 2.6 | 18.3 | 22.2 | 28.0 | 12.6 | 15.8 | 21.0 | 3.2 | 4.4 | 7.2 | 3.5 | 2.3 | 4.4 |
| rec-dom | 2.0 | 27.4 | 31.8 | 39.2 | 19.1 | 22.2 | 26.4 | 5.5 | 7.8 | 12.9 | 6.9 | 8.5 | 5.0 |
| add-add | 1.4 | 6.7 | 7.7 | 10.5 | 47.5 | 51.9 | 55.8 | 2.7 | 3.1 | 4.7 | 7.9 | 9.2 | 6.1 |
| add-dom | 1.7 | 14.2 | 15.9 | 19.0 | 51.6 | 55.7 | 59.2 | 6.0 | 7.5 | 10.3 | 11.1 | 13.3 | 5.4 |
| dom-dom | 1.8 | 7.8 | 9.8 | 14.3 | 63.8 | 66.9 | 70.6 | 2.4 | 3.7 | 7.3 | 14.7 | 17.6 | 6.2 |
| X-rec | 12.2/4.8 | 10.3 | 14.0 | 26.2 | 10.0 | 12.7 | 18.8 | 0.1 | 1.3 | 4.9 | 5.6 | 9.9 | 4.8 |
| X-add | 9.1/2.0 | 33.9 | 39.1 | 48.5 | 24.3 | 25.6 | 31.0 | 3.8 | 5.3 | 11.4 | 21.9 | 35.7 | 5.7 |
| X-dom | 9.8/2.0 | 46.5 | 51.3 | 59.7 | 26.9 | 28.5 | 32.6 | 5.9 | 8.6 | 14.4 | 31.0 | 52.8 | 3.8 |
| FDR <0.1 |  |  |  |  |  |  |  |  |  |  |  |  |  |
| rec-rec | 10.0 | 16.6 | 21.4 | 34.7 | 18.5 | 23.5 | 35.5 | 3.5 | 5.4 | 15.0 | 5.1 | 8.3 | 34.7 |
| rec-add | 5.5 | 32.7 | 39.7 | 52.7 | 27.2 | 32.6 | 45.2 | 11.4 | 15.5 | 27.9 | 13.2 | 18.9 | 32.9 |
| rec-dom | 4.1 | 42.2 | 49.7 | 62.9 | 33.5 | 37.2 | 48.4 | 16.5 | 21.3 | 33.8 | 22.2 | 30.1 | 28.7 |
| add-add | 3.6 | 14.4 | 17.6 | 30.6 | 63.3 | 69.3 | 77.6 | 8.4 | 11.3 | 23.3 | 21.6 | 28.8 | 36.8 |
| add-dom | 3.5 | 26.5 | 31.1 | 42.0 | 65.5 | 70.6 | 78.1 | 18.2 | 22.8 | 33.5 | 29.2 | 39.3 | 29.1 |
| dom-dom | 3.6 | 16.4 | 22.1 | 35.3 | 76.8 | 79.8 | 85.9 | 9.4 | 15.1 | 29.3 | 35.5 | 48.0 | 26.5 |
| X-rec | 12.2/7.8 | 10.3 | 14.0 | 26.2 | 20.0 | 25.2 | 40.5 | 0.7 | 3.1 | 11.0 | 10.3 | 17.5 | 34.6 |
| X-add | 9.1/4.7 | 33.9 | 39.1 | 48.5 | 33.2 | 36.6 | 48.3 | 6.3 | 9.4 | 20.9 | 28.7 | 46.1 | 30.0 |
| X-dom | 9.8/5.0 | 46.5 | 51.3 | 59.7 | 37.0 | 41.2 | 50.9 | 11.4 | 16.3 | 27.2 | 38.8 | 65.5 | 21.6 |

<sup>1</sup>Architecture abbreviations: add – additive; dom – dominant; rec – recessive<sup>2</sup>Locus 1 for autosomal pairs is *musculus* sterile allele; locus 1 for X-autosomal pairs is X-linked<sup>3</sup>‘detected’ - $\geq 1$ significant SNP within given distance criterion<sup>4</sup>Locus 2 for autosomal pairs has a *domesticus* sterile allele; locus 2 for X-autosomal pairs is autosomal<sup>5</sup>Mean number significant SNPs within distance criterion for either locus<sup>6</sup>Mean number significant SNPs on chromosomes not containing ‘causal’ SNPs

It is important to note that our empirical results suggest that the two-locus models used to simulate phenotypes are overly simplified. We predict that involvement of a sterility locus in multiple incompatibilities would reduce the influence of allele/genotype frequencies of any single partner locus on power.

To estimate the false discovery rate from simulations, we classified significant SNPs not located on the same chromosome as either causative SNP as false positives. Choosing an appropriate distance threshold for false vs. true positives on the same chromosome was not obvious given the distribution of distances to causal SNPs (Figure 4-figure supplement 2). We classified significant SNPs <50 Mb from causative SNPs as true positives and excluded SNPs >50 Mb when calculating FDR. Using permutation-based significance thresholds, the median false positive rate was 0.014 (calculated for simulations with ≥10 SNPs within 50 Mb of either causative locus). These results suggest significant SNPs from the GWAS identified using this more stringent threshold are likely to be true positives. By contrast, the median false positive rate was 0.280 using the FDR<0.1 threshold, indicating this threshold is more permissive than predicted. Thus, there is a substantial chance that SNP associations with relative testis weight and expression PC1 identified using this threshold are spurious and evidence is weak for GWAS regions comprising one SNP significantly associated with a single phenotype.

## DISCUSSION

Genetic mapping of testis weight and testis gene expression in hybrid zone mice implicated multiple autosomal and X-linked loci, and a complex set of interactions between loci. These results provide insight into the genetic architecture of a reproductive barrier between two incipient species in nature.

### Association mapping in natural hybrid populations

The potential to leverage recombination events from generations of intercrossing in hybrid zones to achieve high-resolution genetic mapping of quantitative traits has been recognized for decades (reviewed in Rieseberg and Buerkle 2002)1. Until recently, collection of dense genotype datasets and large sample sizes has not been feasible in natural populations due to logistics and costs. This study demonstrates that loci and genetic interactions contributing to reproductive barrier traits can be identified in a GWAS with a modest sample size. Sample sizes approximating those used for human GWAS are not necessary if the prevalence and genetic architecture of the trait of interest are favorable. In general, epistasis makes genetic mapping more difficult. However, for hybrid defects, dependence of the phenotype on epistasis conversely may facilitate mapping. Despite substantial deleterious effects in hybrids, incompatibility alleles are not subject to negative selection within species and may be at high frequency or fixed within species. Hence, the prevalence of affected individuals in a hybrid zone for epistatic traits may be much higher than for deleterious traits in pure populations (*e.g.* disease in humans).

Combining mapping of multiple sterility-related phenotypes substantially improved power to identify sterility loci. We identified a few loci for each phenotype using stringent significance thresholds based on permutation. In addition, most loci identified using more permissive thresholds showed significant associations with more than one phenotype. Spurious associations are unlikely to be shared across phenotypes, thus evidence from multiple phenotypes provided confidence for contributions of 9 genomic regions to testis weight (on the X and 5 autosomes) and 50 genomic regions to expression PC1 (on the X and 18 autosomes).

The high resolution of mapping in the hybrid zone provides an advantage over laboratory crosses. For example, significant regions identified here (median = 2.1 Mb, regions with defined intervals) are much smaller than sterility QTL identified in F_2_s (35.1 Mb; White et al. 2011). Many GWAS regions contain few enough genes that it will be possible to individually evaluate the potential role of each in future studies to identify causative genes. For example, 8/12 testis-weight regions and 28/50 expression PC1 regions contain 10 or fewer protein-coding genes.

We identified candidate genes with known roles in reproduction in four testis-weight regions and 17 expression PC1 regions (Tables 1–2). However, for the majority of regions (8/12 relative testis weight, 33/50 expression PC1), there are no overlapping/nearby genes previously linked to fertility. It is unlikely that these regions would be prioritized if contained in large QTL intervals. High resolution mapping is possible using mapping resources such as the collaborative cross (Aylor et al. 2011) and heterogeneous stocks (Svenson et al. 2012), but these populations represent a small proportion of genetic diversity in house mice (Yang et al. 2011) and hybrid incompatibility alleles may have been lost during strain production.

### Polymorphism of hybrid male sterility loci

Comparisons of different F_1_ crosses between strains of *domesticus* and *musculus* have shown that hybrid sterility phenotypes and loci depend on the geographic origins of parental strains (Britton-Davidian et al. 2005; Good et al. 2008a), suggesting that most hybrid sterility alleles are segregating as polymorphisms within subspecies. Several of the loci identified in this study of hybrid zone mice are novel, providing additional evidence that sterility alleles are polymorphic within subspecies. However, a majority of loci we identified in natural hybrids are concordant with previously identified sterility QTL (Tables 1–2, Figure 2). This similarity suggests there are common genetic factors underlying hybrid sterility in house mice, although there was no statistical support that genome-wide patterns of overlap with previous studies for testis weight or expression PC1 were non-random (*P* > 0.05, 10,000 permutations).

*Prdm9*, discovered by mapping F_1_ hybrid sterility, is the only characterized hybrid sterility gene in mice (Mihola et al. 2009). None of the GWAS regions identified here overlap *Prdm9* (chromosome 17, 15.7 Mb). However, one expression PC1 region (PC42) is ~4 Mb proximal to *Prdm9*. Reductions in PC1 are observed in individuals that are heterozygous or homozygous for the *domesticus* allele at PC42. This pattern is partially consistent with sterility caused by *Prdm9*, which occurs in heterozygous individuals carrying sterile alleles from *domesticus* (Dzur-Gejdosova et al. 2012; Flachs et al. 2012). We did not find evidence for significant associations between SNPs near *Prdm9* and testis weight; the nearest GWAS region (RTW09) is ~41 Mb distal and low testis-weight is associated with the *musculus* allele.

There is concordance between some of the genetic interactions between loci identified here and interactions identified by mapping sterility phenotypes and testis expression traits in an F_2_ cross between *musculus* and *domesticus* (White et al. 2011; Turner et al. 2014) (Figure 2 - figure supplement 1). Precise overlap between some GWAS regions and interaction regions from F_2_s identifies strong candidates for future studies to identify the causative mechanisms and genes underlying sterility loci. For example, an interaction between chromosome 12 and the central X chromosome (RTW11, PC49) identified for testis weight and expression PC1 overlaps an interaction affecting testis expression in F_2_ hybrids (Turner et al. 2014). The 4.3 Mb interval of overlap among chromosome 12 loci (RTW07, PC29, 32.38 – 41.43 Mb F_2_s) encompasses 12 protein-coding genes, including a gene with a knockout model showing low testis weight and sperm count (*Arl4a*) (Schurmann et al. 2002), and two genes with roles in regulating gene expression (*Meox2, Etv1*).

We compared the positions of GWAS regions to 182 regions (163 autosomal, 19 X-linked) with evidence for epistasis based on a genome-wide analysis of genomic clines in a transect across the house mouse hybrid zone in Bavaria (Janousek et al. 2012), the same region where the progenitors of the mapping population were collected. Five testis-weight regions and 18 expression-PC1 regions overlap candidate regions from the hybrid zone genomic clines analysis (Tables 1–2), however the patterns of overlap were not statistically significant (*P* > 0.05, 10,000 permutations). Future introgression analyses using high-density markers within and around GWAS regions may be useful in identifying causative genes and estimating the contributions of sterility alleles to reduced gene flow.

### Role of the X chromosome

Three GWAS regions associated with testis weight and five expression PC1 regions are located on the X chromosome. The X-chromosomal regions surpass the stringent permutation-based significance threshold, and thus have strong statistical support. These results are consistent with evidence for an important role for the X in hybrid sterility from laboratory crosses between subspecies strains geographically diverse in origin (Guenet et al. 1990; Elliott et al. 2001; Oka et al. 2004; Storchova et al. 2004; Oka et al. 2007; Good et al. 2008a,b; Mihola et al. 2009; White et al. 2012) and evidence for greatly reduced gene flow of X-linked loci across the European hybrid zone (Tucker et al. 1992; Payseur et al. 2004; Macholan et al. 2007; Teeter et al. 2008; 2010). A disproportionately large contribution of the X chromosome is a common feature of reproductive isolation in many taxa, the so-called “large X effect” (Coyne and Orr 1989).

The *musculus* derived X chromosome has been implicated repeatedly in genetic studies of sterility in F_1_ and F_2_ hybrids (reviewed in Good et al. 2008a; White et al. 2011). By contrast, *domesticus* alleles were associated with the sterile pattern for most loci we identified on the X in hybrid zone mice (Tables 1–2). A testis expression-QTL mapping study performed in F_2_s showed that *domesticus* ancestry in the central/distal region of the X was associated with a sterile expression pattern (Turner et al. 2014). Differences between studies might reflect geographic variation in sterility alleles but identification of *domesticus*-sterile X alleles only in generations beyond the F_1_ suggests interactions with recessive autosomal partner loci are essential. The importance of recessive sterility alleles was demonstrated by the discovery of multiple novel recessive loci in an F_2_ mapping study (White et al. 2011). F_1_ hybrids are essentially absent in nature (Teeter et al. 2008; Turner et al. 2012), because the hybrid zone is ≥30 km wide (Boursot et al. 1993) thus pure subspecies individuals rarely encounter each other. Consequently, recessive autosomal loci acting in F_2_ and advanced generation hybrids contribute to the maintenance of reproductive isolation in the hybrid zone and may have played important roles in its establishment.

### Genetic architecture of hybrid sterility

Despite a growing list of sterility loci and genes identified in a variety of animal and plant taxa, there are few cases of Dobzhansky-Muller incompatibilities for which all partner loci are known (Phadnis 2011). Hence, there remain many unanswered questions about the genetic architecture of hybrid defects. For example, how many incompatibilities contribute to reproductive barriers in the early stages of speciation? How many partner loci are involved in incompatibilities? Are these patterns consistent among taxa?

The interactions contributing to sterility phenotypes we mapped in hybrid zone mice reveal several general features of the genetic architecture of hybrid sterility. Most sterility loci interact with more than one partner locus. This pattern is consistent with evidence from studies mapping sterility in F_1_ *musculus*-*domesticus* hybrids (Dzur-Gejdosova et al. 2012) and mapping interactions affecting testis gene expression in F_2_ hybrids (Turner et al. 2014). We did not have sufficient power to map interactions requiring three or more sterility alleles, but interactions between alleles from the same subspecies imply their existence. Loci causing male sterility in *Drosophila pseudoobscura* Bogota-USA hybrids also have multiple interaction partners; seven loci of varying effect size interact to cause sterility (Phadnis 2011). In hybrids between *Drosophila koepferae* and *Drosophila buzzatii*, sterility is associated with many loci of small effect, consistent with a polygenic threshold model (Moran and Fontdevila 2014). These studies suggest biological pathways/networks are often affected by multiple Dobzhansky-Muller interactions; a single pairwise interaction between incompatible alleles disrupts pathway function enough to cause a hybrid defect phenotype but when more incompatible alleles are present, the effects of multiple pairwise interactions are synergistic. Variation in the effect sizes of sterility loci might then reflect variation in the number of networks in which the gene is involved, and the connectedness/centrality of the gene within those networks.

Characteristics of the incompatibility network are important for generating accurate models of the evolution of reproductive isolation. A “snowball effect” – faster-than-linear accumulation of incompatibilities caused by epistasis – is predicted on the basis of the Dobzhansky-Muller model (Orr 1995; Orr and Turelli 2001). Patterns of accumulation of hybrid incompatibilities in *Drosophila* and *Solanum* provide empirical support for the snowball hypothesis (Moyle and Nakazato 2010; Matute et al. 2010). Because most GWAS regions have many interaction partners, our results are not consistent with the assumption of the snowball model that incompatibilities are independent, suggesting network models of incompatibilities (Johnson and Porter 2000; Porter and Johnson 2002; Johnson and Porter 2007; Palmer and Feldman 2009) may be more accurate for understanding the evolution of reproductive barriers in house mice.

Involvement of hybrid sterility loci in interactions with multiple partner loci also has important implications for understanding the maintenance of the hybrid zone. Because deleterious effects of a sterility allele are not dependent on a single partner allele, the marginal effect of each locus and thus visibility to selection is less sensitive to the allele frequencies at any single partner locus in the population.

Identifying and functionally characterizing incompatibility genes is an important goal in understanding speciation, but is unrealistic in most non-model organisms. By contrast, mapping reproductive isolation traits in natural populations to identify the number and location of loci and interactions is feasible. General features of the genetic architecture of hybrid sterility – the number of incompatibilities and number and effect size of interacting loci – are arguably more likely to be shared among organisms than specific hybrid sterility genes. Comparison of these features among taxa may reveal commonalities of the speciation process.

## MATERIALS AND METHODS

### Mapping population

The mapping population includes first-generation lab-bred male offspring of mice captured in the hybrid zone (Bavaria) in 2008 (Turner et al. 2012) (Figure 1 - figure supplement 1). We included 185 mice generated from 63 mating pairs involving 37 unrelated females and 35 unrelated males. Many dams and sires were used in multiple mating pairs, thus our mapping population includes full siblings, half siblings and unrelated individuals. Most mating pairs (53 pairs, 149 offspring) were set up with parents originating from the same or nearby trapping locations. Eleven pairs (36 offspring) include dams and sires originating from more distant trapping locations; phenotypes of these offspring were not reported in (Turner et al. 2012).

### Phenotyping

Males were housed individually after weaning (28 days) to prevent effects of dominance interactions on fertility. We measured combined testis weight and body weight immediately after mice were sacrificed at 9-12 weeks. We calculated relative testis weight (testis weight/body weight) to account for a significant association between testis weight and body weight (Pearson’s correlation = 0.29, *P* = 4.9x10^-5^).

We classify individuals with relative testis weight below the range observed in pure subspecies as showing evidence for sterility (Turner et al. 2012). To confirm that this is an appropriate threshold for inferring hybrid defects, we compared this value to relative testis weights reported previously for offspring from intraspecific and interspecific crosses (Good et al. 2008a). The pure subspecies minimum we observed is substantially lower (>2 standard deviations) than means for males from intraspecific crosses (converted from single relative testis weight: *musculus*^PWK^ x *musculus*^CZECH^ - mean x 10.2, standard deviation = 1.2; *domesticus*^LEWES^ x *domesticus*^WSB^ - mean = 11.0, standard deviation = 1.0) and comparable to (within 1 standard deviation) values observed in F_1_ hybrids from 4/7 interspecific crosses that showed significant reductions (mean plus one standard deviation 4.6 – 9.2mg/g).

### Testis gene expression

We measured gene expression in testes of 179 out of the 185 males from the mapping population. Freshly dissected testes were stored in RNAlater (Qiagen) at 4°overnight, then transferred to −20°until processed. We extracted RNA from 15-20 mg whole testis using Qiagen RNeasy kits, and a Qiagen Tissue Lyser for the homogenization step. We verified quality of RNA samples (RIN > 8) using RNA 6000 Nano kits (Agilent) on a 2100 Bioanalyzer (Agilent).

We used Whole Mouse Genome Microarrays (Agilent) to measure genome-wide expression. This array contains 43,379 probes surveying 22,210 transcripts from 21,326 genes. We labeled, amplified, and hybridized samples to arrays using single-color Quick-Amp Labeling Kits (Agilent), according to manufacturer protocols. We verified the yield (>2 μg) and specific activity (>9.0 pmol Cy3/μg cRNA) of labeling reactions using a NanoDrop ND-1000 UV-VIS Spectrophotometer (NanoDrop, Wilimington, DE, USA). We scanned arrays using a High Resolution Microarray Scanner (Agilent) and processed raw images using Feature Extraction Software (Agilent). Quality control procedures for arrays included visual inspection of raw images and the distribution of non-uniformity outliers to identify large spatial artifacts (*e.g.* caused by buffer leakage or dust particles) and quality control metrics from Feature Extraction protocol GE1_QCMT_Dec08.

We mapped the 41,174 non-control probe sequences from the Whole Mouse Genome Microarray to the mouse reference genome (NCBI37, downloaded March 2011) using BLAT ((Kent 2002); minScore = 55, default settings for all other options). Probes with multiple perfect matches, more than nine imperfect matches, matches to non-coding/intergenic regions only, or matches to more than one gene were excluded. A total of 36,323 probes (covering 19,742 Entrez Genes) were retained.

We preformed preprocessing of microarray data using the R package Agi4x44PreProcess (Lopez-Romero 2009). We used the background signal computed in Feature Extraction, which incorporates a local background measurement and a spatial de-trending surface value. We used the “half” setting in Agi4x44PreProcess, which sets intensities below 0.5 to 0.5 following background subtraction, and adds an offset value of 50. Flags from Feature Extraction were used to filter probes during preprocessing (wellaboveBG=TRUE, isfound=TRUE, wellaboveNEG=TRUE). We retained probes with signal above background for at least 10% of samples. We used quantile normalization to normalize signal between arrays. Expression data were deposited in Gene Expression Omnibus as project GSE61417.

To identify major axes of variation in testis expression, we performed a principal components analysis using *prcomp* in R (R Development Core Team 2010) with scaled variables.

### Genotyping

We extracted DNA from liver, spleen, or ear samples using salt extraction or DNeasy kits (Qiagen, Hilden, Germany). Males from the mapping population were genotyped using Mouse Diversity Genotyping Arrays (Affymetrix, Santa Clara, CA) by Atlas Biolabs (Berlin, Germany).

We called genotypes at 584,729 SNPs using *apt-probeset-genotype* (Affymetrix) and standard settings. We used the *MouseDivGeno* algorithm to identify variable intensity oligonucleotides (VINOs) (Yang et al. 2011); 53,148 VINOs were removed from the dataset. In addition, we removed 18,120 SNPs with heterozygosity > 0.9 in any population because these SNPs likely represent additional VINOs. We performed additional filtering steps on SNPs included in the dataset used for mapping. We only included SNPs with a minor allele frequency >5% in the mapping population. SNPs without a genome position or with missing data for >15% of the individuals in the mapping population or pure subspecies reference panel were removed. We pruned the dataset based on linkage disequilibrium (LD) to reduce the number of tests performed. LD pruning was performed in PLINK (Purcell et al. 2007; Purcell n.d.) using a sliding window approach (30 SNPs window size, 5 SNPs step size) and a VIF threshold of 1 x 10^-6^ (VIF = 1/(1-R^2^) where R^2^ is the multiple correlation coefficient for a SNP regressed on all other SNPs simultaneously). This procedure essentially removed SNPs in perfect LD. These filtering steps yielded 156,204 SNPs.

### Ancestry inference

To identify ancestry-informative SNPs, we compared genotypes from 21 pure *M. m. domesticus* individuals (11 from Massif Central, France and 10 from Cologne/Bonn, Germany) and 22 *M. m. musculus* individuals (11 from Námest nad Oslavou, Czech Republic and 11 from Almaty, Kazakhstan) (Staubach et al. 2012).

We used *Structure* (Pritchard et al. 2000; Falush et al. 2003) to graphically represent the genetic composition of our mapping population (Figure 1-figure supplement 1). We included one diagnostic SNP per 20 cM, 3 – 5 markers/chromosome totaling 60 SNPs genome wide. We used the ‘admix’ model in *Structure* and assumed two ancestral populations.

### Association mapping

To identify genomic regions significantly associated with relative testis weight and testis gene expression, we used a mixed model approach to test for single SNP associations. Admixture mapping – often applied in studies using samples with genetic ancestry from two distinct populations – was not appropriate for this study because it was not possible to account for relatedness among individuals in the mapping population (Buerkle and Lexer 2008) (Winkler et al. 2010).

We performed association mapping using GEMMA (Zhou and Stephens 2012), which fits a univariate mixed model, incorporating an *n* x *n* relatedness (identity-by-state) matrix as a random effect to correct for genetic structure in the mapping population. We estimated relatedness among the individuals in the mapping population in GEMMA using all markers and the –gk 1 option, which generates a centered relatedness matrix. For each single SNP association test we recorded the Wald test *P* value. Phenotypes tested include relative testis weight (testis weight/body weight, RTW), testis expression principal component 1 (PC1, 14.6% variance, associated with fertility, Figure 1 - figure supplement 2), and normal quantile ranks of gene expression values for individual transcripts. Neither RTW nor expression PC1 were significantly correlated with age at phenotyping (RTW - cor= −0.02, *P*=0.72; PC1 – cor= 0.01, *P*=0.90), thus we did not include age in the model. SNP data, phenotypic data and kinship matrix to run GEMMA area available through Dryad at: doi:10.5061/dryad.2br40.

To account for multiple testing, we first determined stringent significance thresholds by permutation. We randomized phenotypes among individuals 10,000 times, recording the lowest *P* value on the X and autosomes for each permutation. Thresholds set to the 5th percentile across permutations for RTW were 5.73 x 10^-7^ (autosomes) and 5.83 x 10^-5^ (X chromosome); thresholds for expression PC1 were 1.66 x 10^-8^ (autosomes) and 1.01 x 10^-5^ (X chromosome). Next, we identified regions using a more permissive significance threshold based on the 10% false discovery rate (Benjamini and Hochberg 1995), equivalent to *P* = 3.49 x 10^-5^ for RTW and *P* = 2.86 x 10^-4^ for expression PC1.

To estimate the genomic interval represented by each significant LD-filtered SNP, we report significant regions defined by the most distant flanking SNPs in the full dataset showing *r^2^* > 0.9 (genotypic LD, measured in PLINK) with each significant SNP. We combined significant regions < 10 Mb apart into a single region.

### Testing for genetic interactions

Identifying genetic interactions using GWAS is computationally and statistically challenging. To improve power, we reduced the number of tests performed by testing for interactions only among significant SNPs (FDR < 0.1) identified using GEMMA. We tested all pairs of significant SNPs located on different chromosomes for each phenotype (692 pairs RTW, 82,428 pairs expression PC1). To account for relatedness among individuals we used a mixed model approach, similar to the model implemented in GEMMA. We used the *lmekin* function from the *coxme* R package (Therneau 2012) to fit linear mixed models including the identity-by-state kinship matrix as a random covariate. We report interactions as significant for SNP pairs with *P*<0.05 and FDR<0.1 for interaction terms (RTW: FDR<0.1 ~ *P*<0.02; expression PC1: FDR<0.09 ~ *P*<0.05).

### Mapping Simulations

We performed simulations to evaluate the performance of our mapping approach under varying genetic architectures and allele frequencies. We simulated phenotypes using several genetic models of two-locus epistasis and parameters based on the empirical distribution of relative testis weight. The simulation procedure is illustrated in Figure 4 - figure supplement 1. To preserve genetic structure, we simulated phenotypes using two-locus genotypes from the SNP dataset.

We tested 100 autosomal-autosomal SNP pairs (SNPs on different chromosomes) and 100 X-Autosomal pairs (50 with *domesticus* X-linked sterile alleles and 50 with *musculus* X-linked sterile alleles). The criteria used for choosing ‘causative’ SNPs were a minor allele frequency > 0.05 in the mapping population and fixed in at least one pure subspecies. The ‘sterile’ allele could be polymorphic or fixed within subspecies but the alternate ‘non-sterile’ allele had to be fixed within the other subspecies –e.g. *domesticus* sterile alleles have frequencies 0.05-1.0 in the *domesticus* reference populations from France and Germany and the alternate allele at those SNPs are fixed in *musculus* samples from the Czech Republic and Kazakhstan. For each pair, the ‘causative’ SNPs were randomly selected from all SNPs meeting those criteria (144,506 possible *domesticus* sterile, 124,390 possible *musculus* sterile).

For each SNP pair, we modeled all possible combinations of recessive, additive, and dominant autosomal sterile alleles. For each model type, we assigned mean Z scores for each possible two-locus genotype (Figure 4 – Source data 1). The magnitude of the most severe phenotype (−2.3 standard deviations) is based on observed relative testis weights in the most severely affected males. The mean Z score for heterozygotes in additive models was −1.15. Mean Z scores for non-sterile genotypes in the models were randomly drawn from a uniform distribution between −0.5 and 0.5.

For each SNP pair/architecture, 100 data sets were generated by drawing phenotypes (Z scores) for each individual from a normal distribution with the appropriate two-locus mean and standard deviation = 0.75. The standard deviation value, equivalent to 2.98 mg/g, was chosen on the basis of standard deviations in pure subspecies samples from the mapping population (*domesticus =* 2.13, *musculus* = 3.65; (Turner et al. 2012)). This value is higher than standard deviations in intraspecific F1 males (*domesticus*^LEWES^ x *domesticus*^WSB^ = 1.2, *musculus*^PWK^ x *musculus*^CZECH^ = 1.0; (Good et al. 2008a)), suggesting estimates of mapping power may be conservative.

In total, 90,000 simulations were performed, (9 architectures x 100 SNP pairs x 100 data sets). We identified significant SNPs for each data set using GEMMA, as described above for the empirical data. Testing all pairwise interactions between significant SNPs for each simulated dataset was not feasible computationally. For each SNP pair and architecture, we tested all pairwise interactions for one randomly chosen replicate with 50 – 500 significant SNPs, a range encompassing the observed number of significant associations for relative testis weight and expression PC1. Of the 900 SNP pair/architecture combinations, 889 had at least one replicate with results in this range.

### Significance of overlap between candidate sterility loci

We used permutations to test for non-random co-localization of candidate sterility loci from this study and previous QTL and hybrid zone studies. The locations of significant GWAS regions for relative testis weight and expression PC1 were randomized 10,000 times using BEDTools (Quinlan and Hall 2010). To assess overlap between significant regions for the two phenotypes, we counted the number of RTW regions overlapping PC1 regions (and vice versa) for each permutation. To test for overlap between GWAS identified regions and previously reported candidate regions for related phenotypes, we counted the number of permuted regions overlapping the positions of the published regions (fixed) for each replicate. GWAS regions for both phenotypes were compared to genomic regions with evidence for epistasis and reduced introgression in the Bavarian transect of the hybrid zone (Janousek et al. 2012). In addition, RTW regions were compared to testis weight QTL from mapping studies in F_2_ and backcross hybrids from crosses between subspecies (Storchova et al. 2004; Good et al. 2008b; White et al. 2011; Dzur-Gejdosova et al. 2012) and expression PC1 regions were compared to *trans* eQTL hotspots identified in F_2_ hybrids (Turner et al. 2014).

### Gene annotation

We used ENSEMBL (version 66, February 2012) Biomart to download gene annotations for genomic regions significantly associated with relative testis weight. We identified candidate genes in significant regions with roles in male reproduction using reviews of male fertility (Matzuk and Lamb 2008), manual searches, MouseMine searches for terms related to male fertility (http://www.mousemine.org/) and gene ontology (GO) terms related to male reproduction or gene regulation (plus children): meiosis GO:0007126; DNA methylation GO:0006306; regulation of gene expression GO:0010468; transcription GO:0006351; spermatogenesis GO:0007283; male gamete generation GO:0048232; gamete generation GO:0007276; meiotic cell cycle GO:0051321. Many genes with roles in reproduction reported in publications were not annotated with related GO terms, highlighting the limitations of gene ontology. Moreover, genes causing sterility might not have functions obviously related to reproduction.

## Acknowledgments

We thank Xiang Zhou for expert support using *GEMMA*. We thank Bret Payseur for useful discussion and Diethard Tautz, Luisa Pallares, Trevor Price, Detlef Weigel, Jiri Forejt, Gil McVean and an anonymous reviewer for comments on the manuscript. LMT was supported by postdoctoral funding from the Max Planck Society (to D. Tautz) and by a National Human Genome Research Institute (NHGRI) training grant in Genomic Sciences to the University of Wisconsin (NHGRI 5T32HG002760). Research funding for this project was provided by the Max Planck Society (to D. Tautz) and the Deutsche Forschungsgemeinschaft (SFB-680 to BH).

SNP genotype data were deposited in doi:10.5061/dryad.2br40 and gene expression data in Gene Expression Omnibus GSE61417.

---

**Table 1 – Source data 1.** Protein-coding genes in significant relative testis weight regions.

**Table 2 – Source data 1.** Protein-coding genes in significant testis expression PC1 regions.

**Figure 1 - Figure supplement 1.** Mapping population. (**A**) Location of sampling area (black box) in European house mouse hybrid zone. (**B**) Sampling locations for parents of mice in the mapping population. (**C**) Structure analysis of mapping population. Individuals (vertical bands) are arranged by geographic origin and average percentage alleles from *Mus musculus musculus*.

**Figure 1 - Figure supplement 2.** Principal components analysis of genome-wide gene expression in testis. (**A**) Plot of principal component 1 (PC1) vs. PC2 scores. Individuals with relative testis weight and/or sperm count below the pure subspecies range are indicated in blue (“low fertility”). Individuals with relative testis weight and sperm count within one standard deviation of the mean in pure subspecies individuals are indicated in red (“fertile range”). (**B**) Plot of relative testis weight vs. PC1 score. Correlation coefficient (Pearson’s) and *P* value are indicated.”). (**C**) Plot of hybrid index (% *musculus* alleles on autosomes) vs. PC2 score. Correlation coefficient (Pearson’s) and *P* value are indicated.

**Figure 2 - Figure supplement 1.** Genetic interactions associated with hybrid sterility in hybrid zone mice and in F_2_ hybrids. Orange boxes in outer rings indicate QTL for testis-related phenotypes (testis weight and seminiferous tubule area) identified in previous studies, yellow boxes indicate QTL for other sterility phenotypes and red boxes indicate *trans* eQTL hotspots (see Table 2 for details). Green boxes indicate significant GWAS regions for relative testis weight. Purple boxes indicate significant GWAS regions for testis expression PC1. Lines represent significant genetic interactions identified in hybrid zone mice for relative testis weight (in green) and expression PC1 (in purple) which are concordant with genetic interactions identified by mapping expression traits in F2 hybrids (Turner et al. 2014). Plot generated using circos (Krzywinski et al. 2009).

**Figure 4 - Figure Supplement 1.** Mapping simulation methods. Schematics of (**A**) choice of ‘causal’ SNP pairs from the genotype data, (**B**) phenotype distributions for simulations, (**C**) generation of simulated phenotype data sets, (**D**) association mapping. In (**B**), histogram shows the empirical distribution of relative testis weight in the mapping population, in standard deviation units.

**Figure 4 - Figure Supplement 2.** Distances of significant SNPs to causal SNP in simulations. Distributions are shown at two scales for autosomal and X-linked loci.

**Figure 1 - Source data 1.** SNPs significantly associated with relative testis weight and/or testis expression PC1 (excel file).

**Figure 2 - Source data 1.** Significant genetic interactions (SNP pairs) for relative testis weight (excel file).

**Figure 2 - Source data 2.** Significant genetic interactions (SNP pairs) for testis expression PC1 (excel file).

**Figure 4 - Source data 1.** Z scores for simulation models.

## References

Albrechtová, J., T. Albrecht, S. J. Baird, M. Macholan, G. Rudolfsen, P. Munclinger, P. K. Tucker, and J. Pialek. 2012. Sperm-related phenotypes implicated in both maintenance and breakdown of a natural species barrier in the house mouse. P Roy Soc B-Biol Sci 279:4803–4810. The Royal Society.

Aylor, D. L., W. Valdar, W. Foulds-Mathes, R. J. Buus, R. A. Verdugo, R. S. Baric, M. T. Ferris, J. A. Frelinger, M. Heise, M. B. Frieman, L. E. Gralinski, T. A. Bell, J. D. Didion, K. Hua, D. L. Nehrenberg, C. L. Powell, J. Steigerwalt, Y. Xie, S. N. P. Kelada, F. S. Collins, I. V. Yang, D. A. Schwartz, L. A. Branstetter, E. J. Chesler, D. R. Miller, J. Spence, E. Y. Liu, L. McMillan, A. Sarkar, J. Wang, W. Wang, Q. Zhang, K. W. Broman, R. Korstanje, C. Durrant, R. Mott, F. A. Iraqi, D. Pomp, D. Threadgill, F. Pardo-Manuel de Villena, and G. A. Churchill. 2011. Genetic analysis of complex traits in the emerging Collaborative Cross. Genome Res. 21:1213–1222.

Bateson, W. 1909. Heredity and variation in modern lights. *in* A. C. Seward, ed. Darwin and modern science. Cambridge University Press, Cambridge.

Benjamini, Y., and Y. Hochberg. 1995. Controlling the false discovery rate: a practical and powerful approach to multiple testing. J. Roy. Stat. Soc. Ser. B. (Stat. Method.) 57:289–300.

Boursot, P., J. C. Auffray, J. Britton-Davidian, and F. Bonhomme. 1993. The evolution of house mice. Annu. Rev. Ecol. Syst. 24:119–152.

Briscoe, D., J. C. Stephens, and S. J. Obrien. 1994. Linkage Disequilibrium In Admixed Populations - Applications In Gene-Mapping. J. Hered. 85:59–63.

Britton-Davidian, J., F. Fel-Clair, J. Lopez, P. Alibert, and P. Boursot. 2005. Postzygotic isolation between the two European subspecies of the house mouse: estimates from fertility patterns in wild and laboratory-bred hybrids. Biol. J. Linn. Soc. 84:379–393.

Buerkle, C. A., and C. Lexer. 2008. Admixture as the basis for genetic mapping. Trends Ecol. Evol. 23:686–694.

Carneiro, M., F. W. Albert, S. Afonso, R. J. Pereira, H. Burbano, R. Campos, J. Melo-Ferreira, J. A. Blanco-Aguiar, R. Villafuerte, M. W. Nachman, J. M. Good, and N. Ferrand. 2014. The Genomic Architecture of Population Divergence between Subspecies of the European Rabbit. PLoS Genet. 10:e1003519.

Coyne, J. A., and H. A. Orr. 2004. Speciation. Sinauer Associates, Sunderland, Mass.

Coyne, J. A., and H. A. Orr. 1989. Two rules of speciation. *in* D. Otte and J. A. Endler, eds. Speciation and its consequences. Sinauer Associates, Sunderland, Mass.

Cruickshank, T. E., and M. W. Hahn. 2014. Reanalysis suggests that genomic islands of speciation are due to reduced diversity, not reduced gene flow. Mol. Ecol. 23:3133–3157.

Dobzhansky, T. 1937. Genetics and the origin of species. Columbia University Press, New York.

Dzur-Gejdosova, M., P. Simecek, S. Gregorova, T. Bhattacharyya, and J. Forejt. 2012. Dissecting the genetic architecture of F_1_ hybrid sterility in house mice. Evolution 66:3321–3335.

Ellegren, H., L. Smeds, R. Burri, P. I. Olason, N. Backström, T. Kawakami, A. Künstner, H. Mäkinen, K. Nadachowska-Brzyska, A. Qvarnström, S. Uebbing, and J. B. W. Wolf. 2013. The genomic landscape of species divergence in Ficedula flycatchers. Nature 491:756–760. Nature Publishing Group.

Elliott, R. W., D. R. Miller, R. S. Pearsall, C. Hohman, Y. K. Zhang, D. Poslinski, D. A. Tabaczynski, and V. M. Chapman. 2001. Genetic analysis of testis weight and fertility in an interspecies hybrid congenic strain for Chromosome X. Mamm. Genome 12:45–51.

Falush, D., M. Stephens, and J. K. Pritchard. 2003. Inference of population structure using multilocus genotype data: Linked loci and correlated allele frequencies. Genetics 164:1567–1587.

Filiault, D. L., and J. N. Maloof. 2012. A Genome-wide association study identifies variants underlying the *Arabidopsis thaliana* shade avoidance response. PLoS Genet. 8:e1002589.

Flachs, P., O. Mihola, P. Simecek, S. Gregorova, J. C. Schimenti, Y. Matsui, F. Baudat, B. de Massy, J. Pialek, J. Forejt, and Z. Trachtulec. 2012. Interallelic and intergenic incompatibilities of the *Prdm9* (*Hst1*) gene in mouse hybrid sterility. PLoS Genet. 8:e1003044.

Geraldes, A., P. Basset, B. Gibson, K. L. Smith, B. Harr, H.-T. Yu, N. Bulatova, Y. Ziv, and M. W. Nachman. 2008. Inferring the history of speciation in house mice from autosomal, X-linked, Y-linked and mitochondrial genes. Mol. Ecol. 17:5349–5363.

Good, J. M., M. A. Handel, and M. W. Nachman. 2008a. Asymmetry and polymorphism of hybrid male sterility during the early stages of speciation in house mice. Evolution 62:50–65.

Good, J. M., M. D. Dean, and M. W. Nachman. 2008b. A complex genetic basis to X-linked hybrid male sterility between two species of house mice. Genetics 179:2213–2228.

Good, J. M., T. Giger, M. D. Dean, and M. W. Nachman. 2010. Widespread over-expression of the X chromosome in sterile F_1_ hybrid mice. PLoS Genet. 6:1–13.

Guenet, J. L., C. Nagamine, D. Simonchazottes, X. Montagutelli, and F. Bonhomme. 1990. Hst-3 - an X-Linked Hybrid Sterility Gene. Genet. Res. 56:163–165.

Harrison, R. G. 1990. Hybrid zones: windows on evolutionary processes. Oxford Surveys on Evol Biol 7:69–128.

Hemmer-Hansen, J., E. E. Nielsen, N. O. Therkildsen, M. I. Taylor, R. Ogden, A. J. Geffen, D. Bekkevold, S. Helyar, C. Pampoulie, T. Johansen, FishPopTrace Consortium, and G. R. Carvalho. 2013. A genomic island linked to ecotype divergence in Atlantic cod. Mol. Ecol. 22:2653–2667.

Hewitt, G. M. 1988. Hybrid zones-natural laboratories for evolutionary studies. Trends Ecol. Evol. 3:158–167.

Janousek, V., L. Wang, K. Luzynski, P. Dufkova, M. Vyskocilova, M. W. Nachman, P. Munclinger, M. Macholan, J. Pialek, and P. K. Tucker. 2012. Genome-wide architecture of reproductive isolation in a naturally occurring hybrid zone between Mus musculus musculusand M. m. domesticus. Mol. Ecol. 21:3032–3047.

Johnson, N. A., and A. H. Porter. 2007. Evolution of branched regulatory genetic pathways: directional selection on pleiotropic loci accelerates developmental system drift. Genetica 129:57–70.

Johnson, N. A., and A. H. Porter. 2000. Rapid speciation via parallel, directional selection on regulatory genetic pathways. J Theor Biol 205:527–542.

Johnston, S. E., J. C. McEwan, N. K. Pickering, J. W. Kijas, D. Beraldi, J. G. Pilkington, J. M. Pemberton, and J. Slate. 2011. Genome-wide association mapping identifies the genetic basis of discrete and quantitative variation in sexual weaponry in a wild sheep population. Mol. Ecol. 20:2555–2566.

Kent, W. J. 2002. BLAT---The BLAST-Like Alignment Tool. Genome Res. 12:656–664.

Kocher, T. D., and R. D. Sage. 1986. Further genetic analyses of a hybrid zone between leopard frogs (Rana pipiens complex) in central Texas. Evolution 21–33. JSTOR.

Krzywinski, M., J. Schein, I. Birol, J. Connors, R. Gascoyne, D. Horsman, S. J. Jones, and M. A. Marra. 2009. Circos: An information aesthetic for comparative genomics. Genome Res. 19:1639–1645.

Laurie, C. C., D. A. Nickerson, A. D. Anderson, B. S. Weir, R. J. Livingston, M. D. Dean, K. L. Smith, E. E. Schadt, and M. W. Nachman. 2007. Linkage disequilibrium in wild mice. PLoS Genet. 3:1487–1495.

Lopez-Romero, P. 2009. Agi4x44PreProcess: PreProcessing of Agilent 4x44 array data.

Macholan, M., P. Munclinger, M. Sugerkova, P. Dufkova, B. Bimova, E. Bozikova, J. Zima, and J. Pialek. 2007. Genetic analysis of autosomal and X-linked markers across a mouse hybrid zone. Evolution 61:746–771.

Magwire, M. M., D. K. Fabian, H. Schweyen, C. Cao, B. Longdon, F. Bayer, and F. M. Jiggins. 2012. Genome-wide association studies reveal a simple genetic basis of resistance to naturally coevolving viruses in *Drosophila melanogaster*. PLoS Genet. 8:e1003057.

Malek, T. B., J. W. Boughman, I. Dworkin, and C. L. Peichel. 2012. Admixture mapping of male nuptial colour and body shape in a recently formed hybrid population of threespine stickleback. Mol. Ecol. 21:5265–5279.

Marchini, J., P. Donnelly, and L. R. Cardon. 2005. Genome-wide strategies for detecting multiple loci that influence complex diseases. Nat. Genet. 37:413–417.

Matute, D. R., I. A. Butler, D. A. Turissini, and J. A. Coyne. 2010. A Test of the Snowball Theory for the Rate of Evolution of Hybrid Incompatibilities. Science 329:1518–1521.

Matzuk, M. M., and D. J. Lamb. 2008. The biology of infertility: research advances and clinical challenges. Nat. Med. 14:1197–1213.

Mihola, O., Z. Trachtulec, C. Vlcek, J. C. Schimenti, and J. Forejt. 2009. A mouse speciation gene encodes a meiotic histone H3 methyltransferase. Science 323:373–375.

Moran, T., and A. Fontdevila. 2014. Genome-Wide Dissection of Hybrid Sterility in Drosophila Confirms a Polygenic Threshold Architecture. J. Hered. 105:381–396.

Moyle, L. C., and T. Nakazato. 2010. Hybrid Incompatibility “Snowballs” Between Solanum Species. Science 329:1521–1523.

Muller, H. J. 1942. Isolating mechanisms, evolution and temperature. Biol. Symp. 6:71–125.

Nadeau, N. J., A. Whibley, R. T. Jones, J. W. Davey, K. K. Dasmahapatra, S. W. Baxter, M. A. Quail, M. Joron, R. H. ffrench-Constant, M. L. Blaxter, J. Mallet, and C. D. Jiggins. 2011. Genomic islands of divergence in hybridizing Heliconius butterflies identified by large-scale targeted sequencing. Philosophical Transactions of the Royal Society B: Biological Sciences 367:343–353.

Noor, M. A. F., and J. L. Feder. 2006. Speciation genetics: evolving approaches. Nat Rev Genet 7:851–861.

Noor, M. A. F., and S. M. Bennett. 2009. Islands of speciation or mirages in the desert? Examining the role of restricted recombination in maintaining species. Heredity 103:439–444. Nature Publishing Group.

Nosil, P., T. L. Parchman, J. L. Feder, and Z. Gompert. 2012. Do highly divergent loci reside in genomic regions affecting reproductive isolation? A test using next-generation sequence data in Timema stick insects. BMC Evol. Biol. 12:1–1. BMC Evolutionary Biology.

Oka, A., A. Mita, N. Sakurai-Yamatani, H. Yamamoto, N. Takagi, T. Takano-Shimizu, K. Toshimori, K. Moriwaki, and T. Shiroishi. 2004. Hybrid breakdown caused by substitution of the X chromosome between two mouse subspecies. Genetics 166:913–924.

Oka, A., T. Aoto, Y. Totsuka, R. Takahashi, M. Ueda, A. Mita, N. Sakurai-Yamatani, H. Yamamoto, S. Kuriki, N. Takagi, K. Moriwaki, and T. Shiroishi. 2007. Disruption of genetic interaction between two autosomal regions and the X chromosome causes reproductive isolation between mouse strains derived from different subspecies. Genetics 175:185–197.

Orr, H. A. 1995. The population genetics of speciation: the evolution of hybrid incompatibilities. Genetics 139:1805–1813. Genetics Soc America.

Orr, H. A. H., and M. M. Turelli. 2001. The evolution of postzygotic isolation: accumulating Dobzhansky-Muller incompatibilities. Evolution 55:1085–1094.

Palmer, M. E., and M. W. Feldman. 2009. Dynamics of hybrid incompatibility in gene networks in a constant environment. Evolution 63:418–431.

Payseur, B. A., J. G. Krenz, and M. W. Nachman. 2004. Differential patterns of introgression across the X chromosome in a hybrid zone between two species of house mice. Evolution 58:2064–2078.

Phadnis, N. 2011. Genetic Architecture of Male Sterility and Segregation Distortion in Drosophila pseudoobscura Bogota-USA Hybrids. Genetics 189:1001–1009.

Poelstra, J. W., N. Vijay, C. M. Bossu, H. Lantz, B. Ryll, I. Muller, V. Baglione, P. Unneberg, M. Wikelski, M. G. Grabherr, and J. B. W. Wolf. 2014. The genomic landscape underlying phenotypic integrity in the face of gene flow in crows. Science 344:1410–1414.

Porter, A. H., and N. A. Johnson. 2002. Speciation despite gene flow when developmental pathways evolve. Evolution 56:2103–2111. Wiley Online Library.

Pritchard, J. K., M. Stephens, and P. Donnelly. 2000. Inference of population structure using multilocus genotype data. Genetics 155:945–959.

Purcell, S., B. Neale, K. Todd-Brown, L. Thomas, M. A. R. Ferreira, D. Bender, J. Maller, P. Sklar, P. I. W. de Bakker, M. J. Daly, and P. C. Sham. 2007. PLINK: A Tool Set for Whole-Genome Association and Population-Based Linkage Analyses. The American Journal of Human Genetics 81:559–575.

Quinlan, A. R., and I. M. Hall. 2010. BEDTools: a flexible suite of utilities for comparing genomic features. Bioinformatics 26:841–842.

R Development Core Team. 2010. R: A Language and Environment for Statistical Computing. R Foundation for Statistical Computing, Vienna, Austria.

Renaut, S., C. J. Grassa, S. Yeaman, B. T. Moyers, Z. Lai, N. C. Kane, J. E. Bowers, J. M. Burke, and L. H. Rieseberg. 2013. Genomic islands of divergence are not affected by geography of speciation in sunflowers. Nature Communications 4:1–8. Nature Publishing Group.

Rieseberg, L. H., and C. A. Buerkle. 2002. Genetic mapping in hybrid zones. Am. Nat. 159 Suppl 3:S36–S50.

Rieseberg, L. H., J. Whitton, and K. Gardner. 1999. Hybrid zones and the genetic architecture of a barrier to gene flow between two sunflower species. Genetics 152:713–727.

Rottscheidt, R., and B. Harr. 2007. Extensive additivity of gene expression differentiates subspecies of the house mouse. Genetics 177:1553–1567.

Sage, R. D., J. B. Whitney, and A. C. Wilson. 1986. Genetic analysis of a hybrid zone between domesticus and musculus mice (Mus musculus complex) - hemoglobin polymorphisms. Curr. Top. Microbiol. Immunol. 127:75–85.

Salcedo, T., A. Geraldes, and M. W. Nachman. 2007. Nucleotide variation in wild and inbred mice. Genetics 177:2277–2291.

Schumer, M., R. Cui, D. L. Powell, R. Dresner, G. G. Rosenthal, and P. Andolfatto. 2014. High-resolution mapping reveals hundreds of genetic incompatibilities in hybridizing fish species. eLife 3.

Schurmann, A., S. Koling, S. Jacobs, P. Saftig, S. Krau, G. Wennemuth, R. Kluge, and H. G. Joost. 2002. Reduced Sperm Count and Normal Fertility in Male Mice with Targeted Disruption of the ADP-Ribosylation Factor-Like 4 (Arl4) Gene. Mol. Cell. Biol. 22:2761–2768.

Staubach, F., A. Lorenc, P. W. Messer, K. Tang, D. A. Petrov, and D. Tautz. 2012. Genome Patterns of Selection and Introgression of Haplotypes in Natural Populations of the House Mouse (Mus musculus). PLoS Genet. 8:e1002891.

Storchova, R., S. Gregorova, D. Buckiova, V. Kyselova, P. Divina, and J. Forejt. 2004. Genetic analysis of X-linked hybrid sterility in the house mouse. Mamm. Genome 15:515–524.

Stranger, B. E., E. A. Stahl, and T. Raj. 2011. Progress and Promise of Genome-Wide Association Studies for Human Complex Trait Genetics. Genetics 187:367–383.

Svenson, K. L., D. M. Gatti, W. Valdar, C. E. Welsh, R. Cheng, E. J. Chesler, A. A. Palmer, L. McMillan, and G. A. Churchill. 2012. High-resolution genetic mapping using the Mouse Diversity outbred population. Genetics 190:437–447.

Szymura, J. M., and N. H. Barton. 1991. The genetic structure of the hybrid zone between the fire-bellied toads Bombina bombina and B. variegata: comparisons between transects and between loci. Evolution 237–261. JSTOR.

Teeter, K. C., B. A. Payseur, L. W. Harris, M. A. Bakewell, L. M. Thibodeau, J. E. O'Brien, J. G. Krenz, M. A. Sans-Fuentes, M. W. Nachman, and P. K. Tucker. 2008. Genome-wide patterns of gene flow across a house mouse hybrid zone. Genome Res. 18:67–76.

Teeter, K. C., L. M. Thibodeau, Z. Gompert, C. A. Buerkle, M. W. Nachman, and P. K. Tucker. 2010. The variable genomic architecture of isolation between hybridizing species of house mouse. Evolution 64:472–485.

Therneau, T. 2012. coxme: Mixed Effects Cox Models.

Tucker, P. K., R. D. Sage, J. Warner, A. C. Wilson, and E. M. Eicher. 1992. Abrupt cline for sex-chromosomes in a hybrid zone between two species of mice. Evolution 46:1146–1163.

Turner, L. M., D. J. Schwahn, and B. Harr. 2012. Reduced male fertility is common but highly variable in form and severity in a natural house mouse hybrid zone. Evolution 66:443–458.

Turner, L. M., M. A. White, D. Tautz, and B. A. Payseur. 2014. Genomic networks of hybrid sterility. PLoS Genet. 10:e1004162.

Turner, T. L., and M. W. Hahn. 2010. Genomic islands of speciation or genomic islands and speciation? Mol. Ecol. 19:848–850. Wiley Online Library.

Turner, T. L., M. W. Hahn, and S. V. Nuzhdin. 2005. Genomic islands of speciation in *Anopheles gambiae*. PLoS Biol. 3:e285.

White, M. A., B. Steffy, T. Wiltshire, and B. A. Payseur. 2011. Genetic dissection of a key reproductive barrier between nascent species of house mice. Genetics 189:289–304.

White, M. A., M. Stubbings, B. L. Dumont, and B. A. Payseur. 2012. Genetics and evolution of hybrid male sterility in house mice. Genetics 191:917–934.

Winkler, C. A., G. W. Nelson, and M. W. Smith. 2010. Admixture mapping comes of age. Annu. Rev. Genom. Human Genet. 11:65–89.

Wolf, J., J. Lindell, and N. Backstrom. 2010. Speciation genetics: current status and evolving approaches. Philosophical Transactions of the Royal Society B: Biological Sciences 365:1717–1733.

Yang, H., J. R. Wang, J. P. Didion, R. J. Buus, T. A. Bell, C. E. Welsh, F. Bonhomme, A. H.-T. Yu, M. W. Nachman, J. Pialek, P. Tucker, P. Boursot, L. McMillan, G. A. Churchill, and F. P.-M. de Villena. 2011. Subspecific origin and haplotype diversity in the laboratory mouse. Nat. Genet. 43:648–655. Nature Publishing Group.

Zhou, X., and M. Stephens. 2012. technical reports. Nat. Genet. 44:821–824. Nature Publishing Group.

